# Interictal discharges spread along local recurrent networks between tubers and surrounding cortex

**DOI:** 10.1101/691170

**Authors:** S Tumpa, R Thornton, M Tisdall, T Baldeweg, KJ Friston, RE Rosch

**Affiliations:** Wellcome Trust Centre for Neuroimaging, UCL Queen Square Institute of Neurology, University College London, London, UK; School of Clinical Medicine, University of Cambridge, Cambridge, UK; Department of Clinical Neurophysiology, Great Ormond Street Hospital for Children NHS Foundation Trust, London, UK; UCL Great Ormond Street Institute of Child Health, University College London, London, UK; Department of Neurosurgery, Great Ormond Street Hospital for Children NHS Foundation Trust, London, UK; MRC Centre for Neurodevelopmental Disorders, King’s College London, London, UK; Department of Bioengineering, University of Pennsylvania, Philadelphia PA, USA

## Abstract

The presence of interictal epileptiform discharges on electroencephalography (EEG) may indicate increased epileptic seizure risk and on invasive EEG are the signature of the irritative zone. In highly epileptogenic lesions – such as cortical tubers in tuberous sclerosis – these discharges can be recorded with intracranial stereotactic EEG as part of the evaluation for epilepsy surgery. Yet the network mechanisms that underwrite the generation and spread of these discharges remain poorly understood, limiting their current diagnostic use.

Here, we investigate the dynamics of interictal epileptiform discharges using a combination of quantitative analysis of invasive EEG recordings and mesoscale neural mass modelling of cortical dynamics. We first characterise spatially organised local dynamics of discharges recorded from 36 separate tubers in 8 patients with tuberous sclerosis. We characterise these dynamics with a set of competing explanatory network models using dynamic causal modelling. Bayesian model comparison of plausible network architectures suggests that the recurrent coupling between neuronal populations within – and adjacent to – the tuber core explains the travelling wave dynamics observed in these patient recordings.

Our results – based on interictal activity – unify competing theories about the pathological organisation of epileptic foci and surrounding cortex in patients with tuberous sclerosis. Coupled oscillator dynamics have previously been used to describe ictal activity, where fast travelling ictal discharges are commonly observed within the recruited seizure network. The interictal data analysed here add the insight that this functional architecture is already established in the interictal state. This links observations of interictal EEG abnormalities directly to pathological network coupling in epilepsy, with possible implications for epilepsy surgery approaches in tuberous sclerosis.

**Significance Statement:** Interictal epileptiform discharges (IEDs) are clinically important markers of an epileptic brain. Here we link local IED spread to network coupling through a combination of clinical recordings in paediatric patients with tuberous sclerosis complex, quantitative EEG analysis of interictal discharges spread, and Bayesian inference on coupled neural mass model parameters. We show that the kinds of interictal discharges seen in our patients require recurrent local network coupling extending beyond the putative seizure focus and that in fact only those recurrent coupled networks can support seizure-like and interictal dynamics when run in simulation. Our findings provide a novel integrated perspective on emergent epileptic dynamics in human patients.

## 1 Introduction

Patients with epilepsy experience recurrent seizures caused by abnormal, hypersynchronous brain activity (Fisher et al., 2005). Most patients achieve seizure control with anti-epileptic drugs (AEDs), but approximately one third continue to have seizures despite treatment (Berg and Rychlik, 2015). For these patients, neurosurgical removal of the putative epileptogenic zone has emerged as an efficacious treatment (Rosenow and Lüders, 2001; Duncan et al., 2016). Some patients have seizures after surgery, suggesting that their epilepsy did not arise from focal abnormal activity alone. In fact, epilepsy is now considered a network disorder with emergent dynamics in an abnormally coupled network (Da Silva et al., 2012).

Tuberous sclerosis complex (TSC) is a genetic multisystem disorder, and a leading cause of epilepsy and autism (Osborne et al., 1991). TSC is associated with lesions in many organs including brain, heart, skin, kidneys, and lungs (Curatolo et al., 2008). Intracranial lesions are predominantly cortical tubers, which are highly epileptogenic and characterised by aberrant neuronal coupling in both tuber and surrounding tissue (perituberal cortex) (Ruppe et al., 2014). Tuber core, perituberal cortex, or both have been purported as possible seizure onset zones in patients with TSC (Major et al., 2009; Ma et al., 2012; Kannan et al., 2016). Identifying the local network mechanisms underlying seizures in these patients is a topic under active debate (Gupta, 2017) with implications for surgical treatment, the outcomes of which are currently still variable (Fallah et al., 2015).

Epilepsy surgery in patients with TSC usually involves intracranial EEG (iEEG) evaluation of seizure networks (Bollo et al., 2008), as scalp EEG and imaging alone can be poorly predictive of outcomes (Fallah et al., 2013). Recording iEEG from stereotactically implanted depth electrodes (SEEG) allows sampling from multiple candidate tubers, but the widespread abnormalities, varied seizure propagation pathways, and spatial sampling limitations make evaluating recordings challenging. Extensive resection of the candidate tubers and surrounding cortex are commonly performed even after detailed investigation (Wang et al., 2014). Thus, improved understanding of tuber-related epileptogenicity may help develop more restrictive resections with better surgical outcomes.

The seizure onset zone – the target of neurosurgical intervention – is defined by the ictal activity. Interictal epileptiform discharges (IEDs) can provide complementary localising information (Staley and Dudek, 2006; Murakami et al., 2016), indicate dynamic changes in seizure risk (Baud et al., 2018) and are associated with synaptic connectivity changes potentially contributing to epileptogenesis (Staley et al., 2005). IEDs with the same features are seen across widely spaced electrodes with latencies of tens of milliseconds, suggesting rapid propagation to distant cerebral structures (Emerson et al., 1995; Alarcon et al., 1997; Wendling et al., 2012)

This propagation speed is in stark contrast to epileptic seizures themselves – ictal recruitment often takes several seconds. Yet the same network supports slow ictal, and fast interictal propagation (Proix et al., 2018). Spatiotemporal dynamics of IEDs may thus reveal additional insights into the functional organisation of cortical areas involved in seizure generation and spread (Bettus et al., 2008).

Here, we quantify and model the dynamics of individual IEDs, to infer a neurobiologically plausible connectivity structure. We identify systematic IED phase delays between neighbouring SEEG channels. We then fit network architectures that recapitulate two possible propagation modes: (i) propagation along recurrent coupled oscillators and (ii) propagation of pulses in an excitable network (Muller et al., 2018; Proix et al., 2018). We hypothesised that the fast spread of interictal discharges may reveal existing recurrent coupling in local epileptogenic networks, which can ultimately support both ictal and interictal discharges.

To compare different explanatory networks, we use Dynamic Causal Modelling (DCM), a framework that allows quantitative comparison of candidate explanatory models (Kiebel et al., 2008; Moran et al., 2008, 2011) using neural mass models of neuronal population dynamics (e.g. the canonical microcircuit (Bastos et al., 2012) used here). Recent advances in the efficient estimation across large model spaces (Friston et al., 2018) furthermore allow us to compare model architectures at the level of individual tubers.

## 2 Materials and Methods

### 2.1 Patient identification and research ethics

We retrospectively considered all patients admitted to Great Ormond Street Hospital for Children between January 2016 and June 2018 and included them if they (i) had a diagnosis of TSC, and (ii) underwent presurgical evaluation with SEEG recording for pharmacoresistant epilepsy. All eight patients had focal seizures, and active focal epilepsy with onset in infancy or early childhood. Patient details are summarised in Table **1**. The retrospective use of anonymised clinical data was approved by the UK Health Regulary Authority (HRA, IRAS ID 229772) and the Great Ormond Street Hospital / UCL Great Ormond Street Institute of Child Health Joint Research Office (Project ID 17NP05).

**Table 1:** Patient details

| Patient | Age at epilepsy onset (y) | Age at SEEG recording (y) | Number of SEEG Electrodes |
| --- | --- | --- | --- |
| TS_01 | 1.50 | 8 | 6 |
| TS_02 | 0.50 | 5 | 8 |
| TS_03 | 1.50 | 5 | 16 |
| TS_04 | 1.00 | 5 | 9 |
| TS_05 | 0.06 | 11 | 12 |
| TS_06 | 0.33 | 10 | 9 |
| TS_07 | 0.67 | 12 | 5 |
| TS_08 | 0.22 | 8 | 7 |

### 2.2 Intracranial EEG acquisition, selection and pre-processing

#### 2.2.1 SEEG recording

SEEG electrodes were placed based on multidisciplinary SEEG planning involving neurologists, neurophysiologists and neurosurgeons according to clinical need, informed by presurgical imaging, seizure semiology, and ictal and interictal scalp EEG. DIXI medical SEEG depth electrodes were inserted under stereotactic guidance using the Neuromate Renishaw robot system (programmed with patient specific pre-operative imaging) into various structures, including multiple tubers in each child (Sharma et al., 2019). Most electrodes had one or more contacts in the tuber core, with additional contacts superficial, and/or deep to the core (i.e., in superficial grey matter, and deep white matter respectively, Figure **1A**). SEEG was recorded continuously for up to 5 days in order to capture seizures for clinical interpretation. All SEEG recordings were recorded using a Natus NeuroWorks system at a sampling rate of 1kHz, with a white matter contact remote from regions involved in the generation of seizures used for reference. Quantitative analysis of phase delays was performed on bipolar montage.

**Figure 1.**
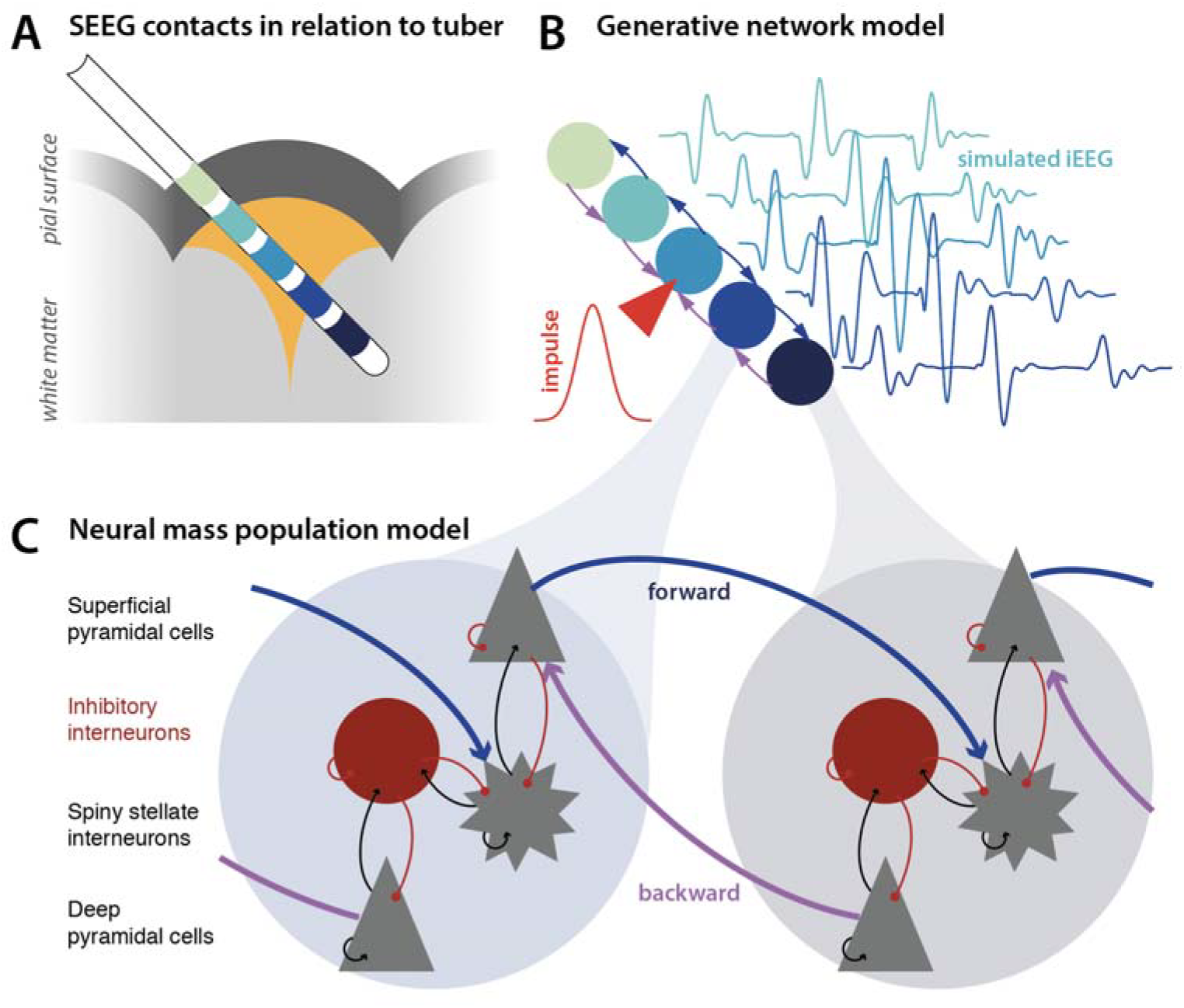
Dynamic causal modelling. (A) Intracranial interictal recordings were acquired with stereotactically implanted depth electrodes (SEEG), and visually classified in terms of their relation to the tuber core. (B) Dynamic causal modelling allows the testing of network models of dynamics that generate iEEG responses to endogenous fluctuations or spikes. We inverted such models to best explain interictal epileptiform discharges. (C) Each node, or ‘source’ in the model comprises coupled neural mass models of 4 populations, organised into two oscillator pairs (superficial pyramidal cells and spiny stellate cells; deep pyramidal cells and inhibitory interneurons).

#### 2.2.2 Classification of SEEG contact position

Pre-implantation MRI FLAIR, T1 and T2-weighted images (WI) were co-registered with post-implantation CT to visually determine the position of SEEG contacts in relation to cortical tubers, perituberal grey, and white matter. In young children, tubers appear hypointense on the T1-WI image and hyperintense on T2-WI and FLAIR images (Grajkowska et al., 2010). Although in some patients, generalised brain volume loss was observed, most affected gyri were enlarged with blurring of the grey-white junction (Jurkiewicz et al., 2006; Weisenfeld et al., 2013). Individual contacts were classified in terms of their position to the tuber core on an integer scale centred around 0 (tuber core position) by consensus of two raters (MT, ST), including a neurosurgeon with clinical expertise in visual identification of TSC features on imaging.

#### 2.2.3 Time series extraction

A total of 15 min of 60-second segments of interictal extra-operative SEEG recordings were selected randomly from the entire recording using a MATLAB implemented random number generator (MATLAB) – segments containing visible artefact, or clinically labelled seizure activity were excluded. Band pass filter (0.5-120Hz, zero phase) and notch filter (50Hz) were applied for visual inspection on the clinical Natus Database system. The signal was re-montaged to bipolar montages between adjacent contacts on the same SEEG electrode and exported in EDF+ format for further processing using custom MATLAB code available online (https://github.com/roschkoenig/Travelling_Spikes).

#### 2.2.4 Spike detection

A custom spike detection algorithm (adapted from SPKDT v1.0.4, (Barkmeier et al., 2012)) was run to identify individual interictal epileptiform discharges. Criteria for spike detection were: (i) peak amplitude >4 standard deviations from the mean, (ii) ascending and descending absolute slope values of >7µ*V/ms*, (iii) total width of spike < 20ms. We then grouped spikes recorded from different contacts of the same SEEG electrode and detected groups of spikes that co-occurred within 200ms of each other in time across channels on the same SEEG electrode. Only spike groups with at least 2 spikes within this time window were considered for further analysis.

### 2.3 Sensor space analysis

In the first instance, we analysed data features of the recorded SEEG signal to determine whether the spatiotemporal distribution of interictal spikes was consistent with a travelling-wave spread and tested these using Bayesian statistical criteria.

#### 2.3.1 Delay estimation

We estimated the temporal delay between clusters of spikes spanning several channels of the same SEEG electrode, by calculating cross correlation between the signal, and estimating the phase-delay between neighbouring channels. This was performed in two frequency bands, aiming to identify phase delays separately for low (<13Hz) and high (13-120Hz) frequency components of the interictal epileptiform discharge.

#### 2.3.2 Bayesian statistics on spatiotemporal spike distribution patterns

To identify the spatiotemporal organisation of interictal discharges, we compared different explanatory models linking the relative time of spike detection to the position of the channel in relation to the tuber core. We proposed three different possible (linear) models that could explain spatiotemporal organisation (where *t* is time, and *p* is position relative to the tuber, with deep white matter represented by the most negative value, the core position represented by zero, and overlying grey matter represented by positive values):

##### Uniform

If there is no spatiotemporal pattern, time of detection should be independent of channel position, modelled through a simple constant function:

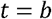

##### Depth to Surface

If interictal discharges spread through the cortical tuber along the grey-white matter axis, we would expect a linear relationship between channel position and time of spike detection as follows:

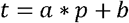

##### Core to Periphery

This model encodes a spread of interictal discharges between core and surrounding tissue through a relationship of absolute distance from core, and time of detection:

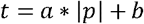

We fit each of these simple linear models to the spike groups detected in the earlier steps and compared models using the Bayesian Information Criterion (BIC) for each of these models. Unlike some other criteria, BIC penalises for additional parameters (i.e. added complexity). Lower BIC values imply either fewer explanatory variables, better fit or both (Schwarz, 1978), and the model with the lowest BIC has the best explanatory power for the observed data under Bayesian constraints. The Bayes factor between competing models can be approximated by the difference in BIC as follows: 2 log_*e*_(*B*_10_) ~2(*ΔBIC*); where a value of >10 is considered very strong evidence (Jones et al., 2001).

### 2.4 Dynamic causal modelling

#### 2.4.1 DCM framework

Macroscopic neuronal dynamics as measured by iEEG can be generated by mesoscale neural mass models that describe the average behaviour of neuronal populations and their coupling, as pioneered by Wilson & Cowan (Wilson and Cowan, 1973). Here, we use the dynamic causal modelling (DCM) (Friston et al., 2003) to compare different models of local network coupling underlying the dynamics of interictal discharges. We use the canonical microcircuit model (Bastos et al., 2012), which is widely used in DCM, including work in epilepsy and other neurological abnormalities (Auksztulewicz and Friston, 2015; Bastos et al., 2015; Litvak et al., 2015; Papadopoulou et al., 2015; Pinotsis et al., 2017). This model comprises four coupled neural masses, organised in two excitatory/inhibitory oscillator pairs, and parameterised by coupling strength parameters and time constants.

DCM assumes that the activity in one source is evoked by the activity in another (David and Friston, 2003), and DCM for EEG uses the above mentioned neural mass model to explain the source activity, and causal interactions (Garrido et al., 2008). Rather than estimating source activity at isolated points in time, it models source activity over time, accounting for the interacting inhibitory and excitatory populations of neurons. DCM uses variational Bayesian model inversions to infer the parameters that best explain the source activity and provides a free energy approximation to the model evidence. This can be used for Bayesian model comparison (c.f., the use of BIC above). To identify the architecture of local perituberal networks, we constructed multi-region DCMs, with each region containing a full canonical microcircuit and located along a deep white matter – tuber core – perituberal grey matter (depth to surface) axis. We compared their model evidence (and accompanying connectivity estimates) as outlined below. In order to accommodate the diversity of possible IED for each tuber included, we performed *k*-means clustering on the loading of the principal components that accounted for at least 90% of the overall variance of IED multichannel time series for each tuber, identifying clusters of IEDs with shared similar morphologies. We then selected the IEDs closest to the centroids of each cluster as representative and included these episodes in the modelling. These data were inverted in a time window of −100ms to 250ms around the peak of the first spike detection with a temporal Hanning window, using a Gaussian impulse input function at 0ms with 16ms SD (to model an endogenous discharge event).

#### 2.4.2 Model Specification

We specified our candidate network models of local coupling along two main dimensions:

1. <u>Recurrent vs outward only coupling</u>: Previous mathematical models of travelling wave propagation and ictal wave propagation suggested different modes of wave propagation (Ermentrout and Kleinfeld, 2001; Smith et al., 2016; Muller et al., 2018): (i) Travelling waves propagating through excitable neural media. Here, the wave originates at a single oscillator (i.e., a pacemaker), directly exciting the neighbouring regions of the cortex through a progression of increasing time delays. (ii) Message passing along a recurrently coupled network. Crucially in this model architecture, all local oscillators are potential sources for rhythmic outputs.
2. <u>Superficial, core, or deep source</u>: In the neural mass formulation, an impulse will trigger the subsequent spread of activity. Here, we model this as Gaussian input into key positions of the network established above – specifically the perituberal gray (most superficial superficial), the tuber core itself (core), or the deep white matter (deepest) nodes.

This gives us a model space of 2*3 = 6 models which we compared using a (variational) free energy bound on Bayesian model evidence (see below).

#### 2.4.3 Bayesian model reduction and model comparison

DCM, through variational Bayesian inference (specifically, Variational Laplace), allows estimation of posterior densities over model parameters, given the data. Model inversion also provides a free energy bound on log model evidence (i.e. *F* ≈ − ln *p*(*y*|*m*)). This free energy can be efficiently approximated for nested (i.e., reduced) models whose parameters are subsets of parameters of a parent (i.e., full) model, by running the full model inversion once – followed by Bayesian model reduction (Friston et al., 2018). The difference in free energy values between competing models corresponds to the logarithm of the corresponding Bayes factor (Baele et al., 2013; Lin and Yin, 2015), which we use here for model comparison. For group level comparison, we used a random effects Bayesian analysis (Penny et al., 2010), accounting for potential individual differences in network architecture between tubers, which we will report as exceedance probability of the competing models.

#### 2.4.4 Simulations

Finally, we took a representative example of a full network fitted in the previous step, and simulated changes in intrinsic excitability, to simulate possible transitions into ictal dynamics (Papadopoulou et al., 2015). Here, the DCM is used in ‘simulation mode’, meaning that instead of inferring parameter values from given data, we simulated data based on fluctuations about empirically determined model parameters (i.e., synaptic connectivity and related time constants). We repeated these simulations with models that did, and did not contain recurrent coupling between neighbouring nodes, to illustrate the relationship between model topology and ensuing dynamics.

## 3 Results

### 3.1 Interictal discharges form travelling waves between tuber core and periphery

Spatially coordinated phase delays (Figure **2**) between spikes on different channels that belong to the same electrode suggest that these interictal discharges behave like a travelling wave.

**Figure 2.**
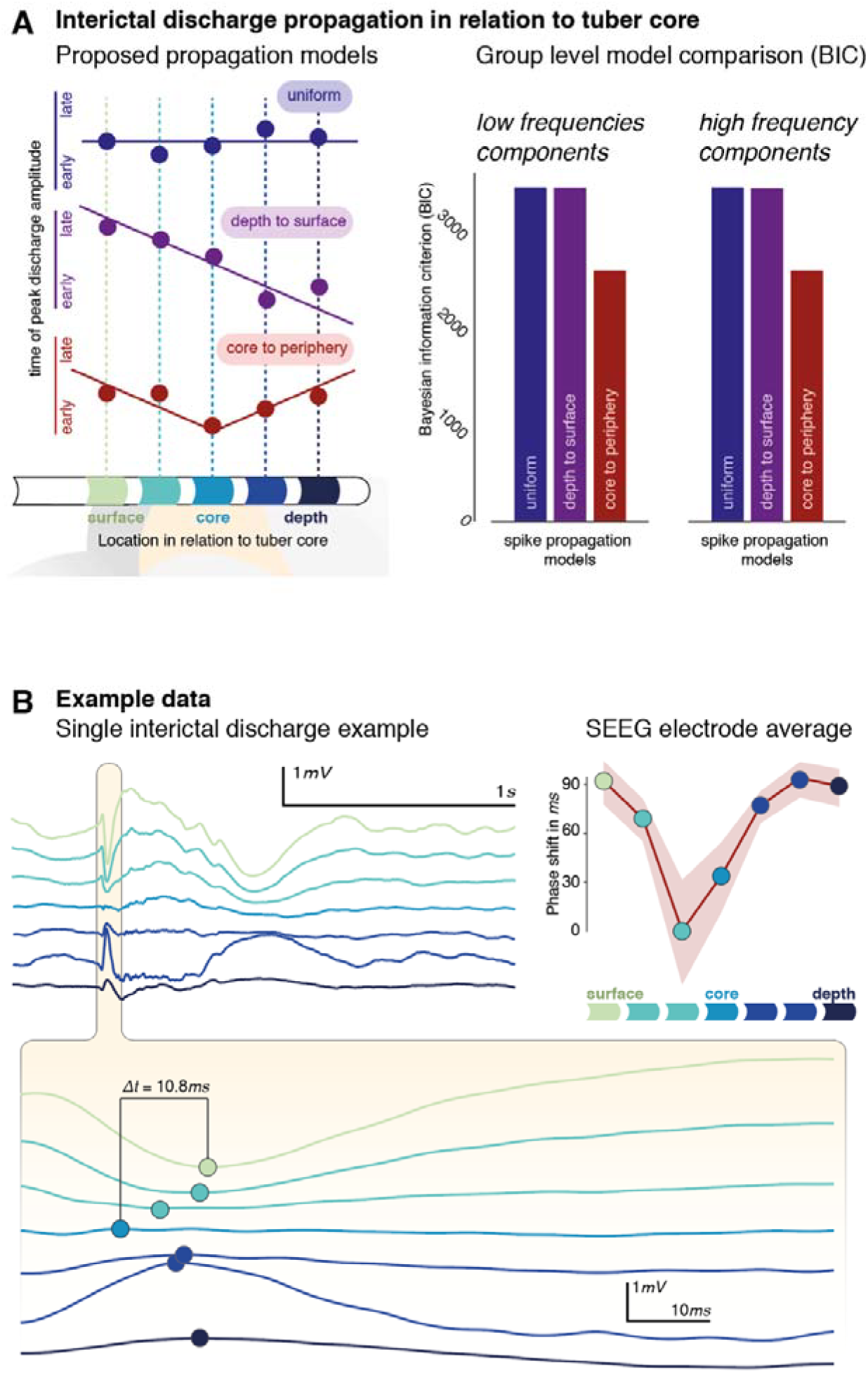
Phase delays follow a tuber core to periphery gradient. (A) Three linear models were used to identify possible relationships between phase delays and spatial location of SEEG contacts. These explain the relationship either as *uniform* (no relationship), *depth to surface*, or *core to periphery* local spread of putative IED travelling waves. Model comparison identifies the *core to periphery* organisation of IEDs as the most likely model (with the lowest BIC value). (B) Travelling wave dynamics can also be identified at the level of individual IEDs, as shown in the core to periphery organisation of temporal differences between spike peak timings in the bottom panel. For the same electrode, the average phase shift between detected groups of IEDs is shown for individual SEEG contacts against their spatial location in relation to the tuber core (shaded area illustrates standard error around the mean).

We used the Bayesian information criterion – on a set of simple linear models – to identify the spatiotemporal dynamics of individual interictal discharges. The linear models characterise the relationship between relative phase delay and SEEG contact position as either (i) *uniform*; i.e. no relationship between timing of spike detection and SEEG contact position, (ii) *depth to surface*; i.e. there is a gradient of delays between the deepest, and the most superficial contact of the SEEG electrode, and (iii) *core to periphery*; i.e. there is a gradient of delays between contacts closest to the tuber core, and contacts peripheral to this (Fig **2A**). Models were estimated separately for low (1-13Hz), and fast (13-120Hz) frequency components to account for potential propagation differences between fast and slow components of the interictal epileptiform discharges.

The winning model at the group level (i.e., the one with the lowest BIC value) is the *depth to surface* model for both frequency bands considered (Figure **2A**). The BIC difference is >800, greatly exceeding the commonly considered ‘significance’ threshold of ΔBIC > 10 (Jones et al., 2001). This characterises IEDs as waves travelling between the tuber core and its periphery. This travel can also be identified on the level of individual spikes, and on electrode-wide averages (Figure **2B**). There was no statistical relationship between propagation patterns at the individual SEEG electrodes and their clinically reported involvement in clinical seizure onset (*X*^2^ (1, N = 38) = 1.59, p=0.207).

### 3.2 Interictal discharges are supported by recurrently coupled local networks

Using Dynamic causal modelling, we compared how plausible local network architectures explain the observed spatiotemporal dynamics of the IEDs. We organised the model space along two main dimensions: (1) the origin of the IED, modelled here as input into one of three network nodes, and (2) two coupling schemes between neighbouring nodes in the network (Figure **3A**). A full model was fitted to a clustering-based selection of up to 3 IEDs per included tuber. The parameters and free energy of the full 3 * 2 = 6 model space were then estimated using Bayesian model reduction. The models were compared at the group level using a random effects analysis based on the model- and subject-specific free energy estimates of the model evidence. This provides an estimate of the exceedance probability that allows for comparison of explanatory models reported here (Figure **3B**).

**Figure 3.**
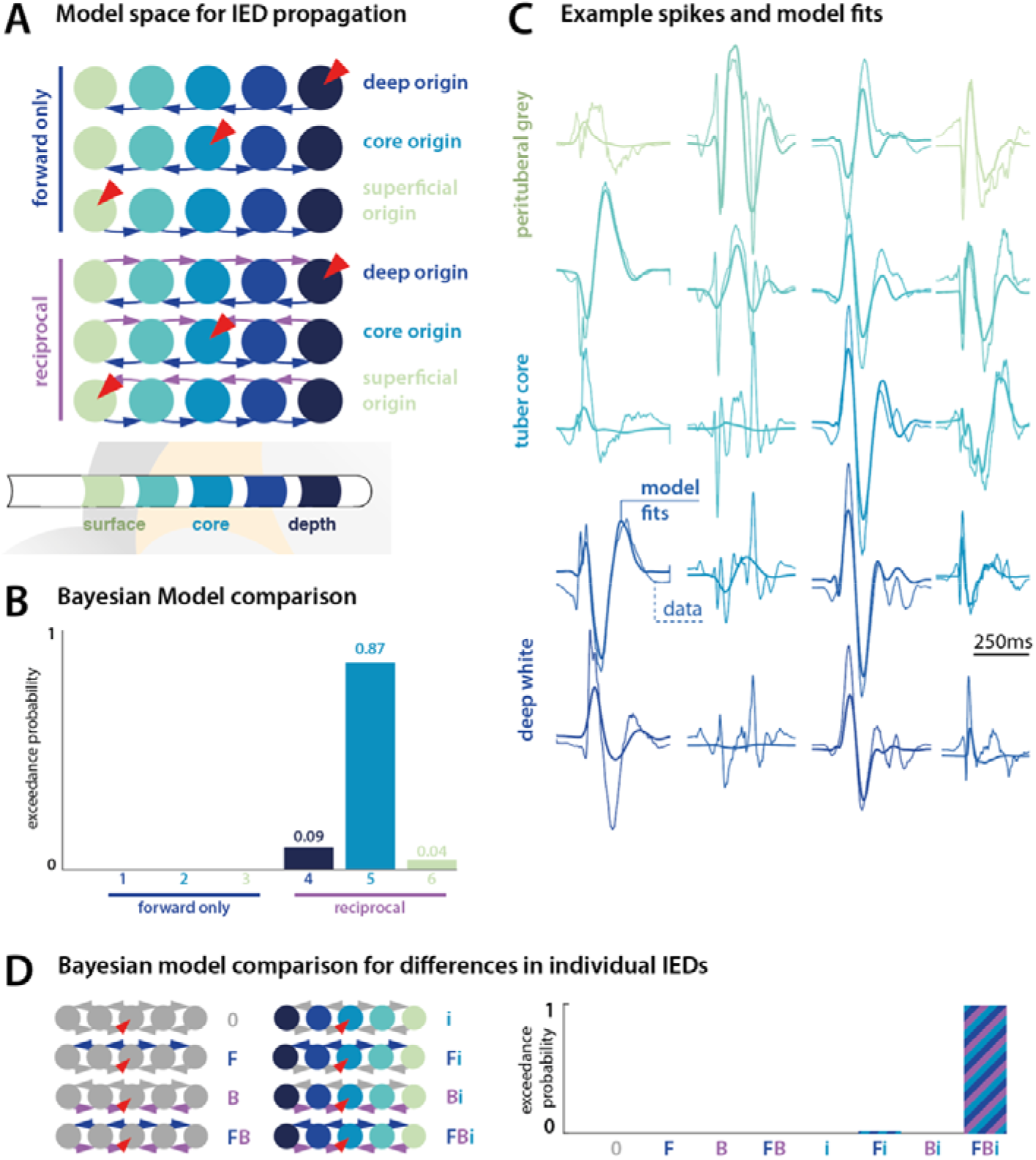
Recurrent coupling in local networks supports IED. (A) Six candidate networks we used to explain IED spread around the tuber – 3 sets of forward coupled, or recurrently coupled networks, each with either superficial, core, or deep origin of the IED. (B) Bayesian model comparison shows that recurrent coupling is necessary to explain IED dynamics, and that across the group spread from the tuber core was the most likely local network architecture. (C) The final winning model captured an average of ~60% of the variance of single spike dynamics across patients, with some representative model fits shown here. (D) In an additional set of models, we explored whether within-electrode differences in individual IEDs can be explained by a subset of models. However, there was strong evidence for a contribution of forward, backward and intrinsic connectivity to observed within-electrode variability in IEDs.

The winning model featured a triggering impulse at the tuber core, with recurrent forward/backward coupling to its neighbouring nodes. This further supports the observation that travelling wave dynamics organise around a core-periphery gradient, but adds the insight that recurrent coupling is required to support the observed IED spread. The winning model explained an average of 58% of the variance for each electrode, with large amplitude, lower frequency wave forms generally captured better (representative examples shown in Figure **3C**).

We further explored whether within-electrode differences in IEDs can be explained by changes in just a few key connections, by comparing different reductions of the winning model architecture. However, to explain the diversity of IEDs observed patients, reciprocal extrinsic connections, and recurrent self-connections were all required (Fig **3D**)

### 3.3 Only recurrently coupled network support both IED- and seizure-like dynamics *in silico*

The network models fitted in the previous step provide us with fully specified generative models, allowing for simulations under different parameter values. Here, we systematically varied the intrinsic excitability (i.e. gain) parameters of neuronal populations for a fully parameterised model from the previous step, both in the recurrent coupling, and the recurrent only coupling network architectures (Fig **4**). These simulations reveal that while fast frequency rhythms akin to ictal discharges can be produced by both recurrent, and forward only network architecture, the IED-like dynamics only emerge in the recurrently coupled network.

**Figure 4.**
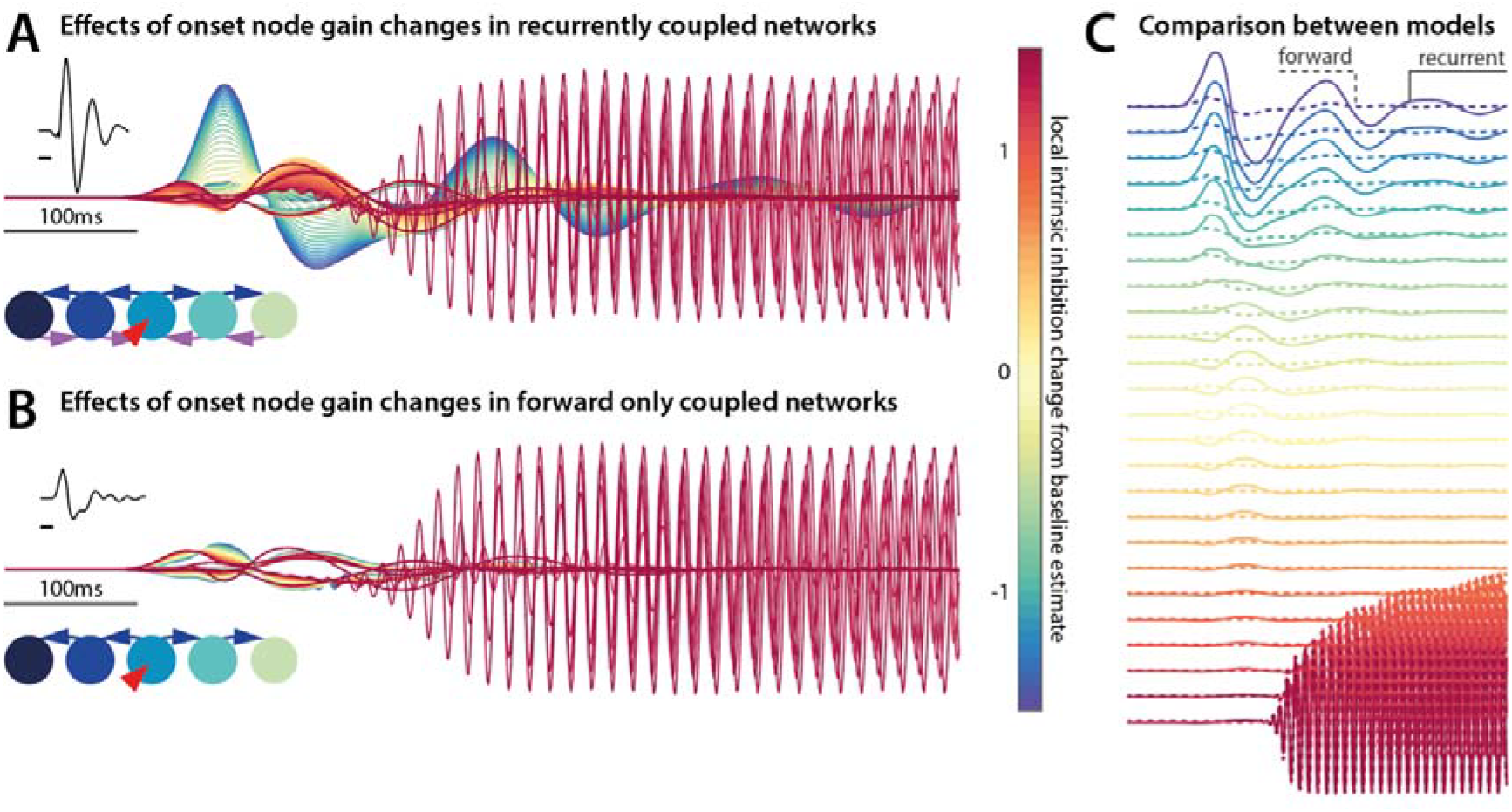
Only networks with recurrent coupling support IED- and seizure-like dynamics. (A) Using a network fitted to patient data as template, we simulate changes in local intrinsic gain of the IED onset node (here the tuber core). Increases in gain allow for fast oscillations akin to seizure-onset rhythms to emerge. (B) Using a model without recurrent coupling reproduces the fast frequency ictal-like discharges, but not the high amplitude synchronous IED-like discharges. (C) The direct comparison between these simulated time series shows the large amplitude difference at low intrinsic gain values.

## 4 Discussion

For this study, we used quantitative analysis and computational modelling to investigate the spatiotemporal dynamics and local network constraints of IEDs in patients undergoing evaluation for epilepsy surgery. We provide three key insights that contribute to current debates in the field: (1) Through quantitative analysis, we identify tuber cores as the spatial source of interictal discharges; (2) With dynamic causal modelling, we show that locally recurrent coupling is necessary to explain the travelling wave dynamics of IEDs; and (3) using simulations we can link these insights on the network constraints to the spatiotemporal dynamics of seizures.

### 4.1 Interictal discharges in tuberous sclerosis spread from the tuber core

This study confirms previous EEG findings that interictal epileptiform discharges (IEDs) form travelling waves and propagate through the cerebral cortex (Emerson et al., 1995; Alarcon et al., 1997). Furthermore, the phase delays of IEDs (in ‘sensor space’, i.e. in the SEEG time series) are organised spatially in relation to the tuber core (Fig 2). This insight was further supported by model comparison of different network origins of the interictal discharge impulse, i.e. in ‘source space’. The model with IEDs starting from the core provides the overall best explanation of the observed dynamics (Fig 3).

In addition to further characterising IEDs generally, the findings also add to the previous literature on ictogenesis of lesions and surrounding grey matter in patients with TSC specifically. Histologically, tuber cores show dysmorphic neurons and balloon cells, which are also found in other epileptogenic lesions such as focal cortical dysplasia type IIb (André et al., 2007). However, dyslamination and cellular dysplasia are also reported to affect the perituberal cortex (Ruppe et al., 2014), providing a possible pathophysiological role for the peritubular cortex in ictogenesis. There has been conflicting evidence from iEEG and corollary investigations suggesting that seizures may either emerge from the core of the tuber (Mohamed et al., 2012; Kannan et al., 2016), or the surrounding cortex (with the tuber core themselves eventually becoming electrographically silent (Major et al., 2009; Ma et al., 2012).

We provide corroborating evidence that IEDs recapitulate a core-to-periphery organisation of abnormal brain dynamics, even outside of seizures. As in many cases of IEDs in patients with focal epilepsy, the relationship between IEDs and ictal dynamics at the individual level is nontrivial: We observed some variability in the direction of IED travel for individual tubers (cf. Fig 3B – where models 4 and 6 had small but identifiable exceedance probabilities), but the direction of local IED spread around individual tubers was not strongly associated with clinical labelling as to whether a given tuber was involved at seizure onset.

To integrate the insights on local IED spread further into a model of local networks surrounding epileptogenic lesions, we used dynamic causal modelling. This allowed us to generate and compare biophysically plausible network models – enabling us to identify the network constraints underlying the observed IED spread.

### 4.2 Recurrent local coupling is necessary for IED dynamics

We used DCM to test competing mechanistic accounts of the propagation of IEDs along several contacts of the SEEG recording. Instead of exploring all the possible models, DCM tests specific models of connectivity and by selecting one, can provide evidence in favour of that model in comparison to plausible alternatives (Kiebel et al., 2008). The networks we examined here were motivated by two complementary imperatives: (a) To explore the onset and propagation of the IEDs in relation to the tuber anatomy, testing whether these evolve in line with previous reports of the ictal waves originating from the tuber core (Kannan et al., 2016); and (b) to identify the organisational principles of networks that support the local spread of IEDs. Phase delays in networks (i.e. travelling waves) can emerge from a number of different architectures (Ermentrout and Kleinfeld, 2001) of which two – forward coupling, vs recurrent coupling – were modelled here.

Within this model space we found evidence for (a) a privileged role of the tuber core as the origin of IEDs, and (b) locally recurrent coupling in the network architecture. This recurrent coupling confers coupled oscillatory dynamics on distributed responses, allowing for arbitrarily fast phase delays of travelling waves to emerge (unlike e.g., the forward-only coupled network). This offers a novel perspective on the functional organisation of the local ictogenic network surrounding the tuber core: Whilst the tubers (particularly in our paediatric sample) are apparently the origin of abnormal activity, it is their close, recurrent integration with surrounding cortex that allows for IED dynamics to emerge. A strongly recurrently coupled network like this may itself support emergent dynamics such as seizures, even if a possible initial ‘pacemaker’ is eventually removed (e.g. through eventual calcification of the tuber core, or focal neurosurgical lesioning).

### 4.3 Coupled-oscillator dynamics support both ictal- and IED dynamics

The models derived from the DCM analysis are not only descriptions of the observed data, but fully generative models, constrained by empirical observations. This means that through simulations, we can explore unobserved parameter combinations, and identify how different parameters generate neuronal dynamics. Here, we used a DCM – with empirically optimised parameters – to reproduce (*in silico*) a transition from IED dynamics to fast oscillation onset rhythm, akin to seizure onset-like dynamics (Fig **4**). We directly compared the effects of increasing intrinsic gain in the network nodes in the context of a network with recurrent coupling and an architecture with forward only coupling. These simulations suggest that only the recurrent coupling architecture can support both near synchronous, large amplitude, low frequency IED like discharges, as well as the high frequency rhythms that characterise seizure-onset dynamics. Crucially, the forward only coupling network did not generate IED like dynamics but was still able to generate the fast seizure-like rhythms.

This simulation illustrates the fact that two dynamical regimes can emerge in a single seizure: whilst the ictal wave front is often characterised by fast frequency rhythms – that slowly recruit more cortical areas over time – ictal spikes later in an established seizure can propagate at near synchronous speeds in the areas of cortex that have already been recruited (Smith et al., 2016; Schevon et al., 2019). These different types of propagation mechanisms can be described in different models, such as propagation in excitable media (ictal onset), or coupled oscillator dynamics (for ictal spike and wave discharges).

Recent computational modelling work has shown that variations in slowly evolving states within a single heterogeneously coupled network of neural masses can support both types of dynamics by transitioning through different types of dynamical regimes during a modelled seizure (Proix et al., 2018). The work presented here suggests – in addition – that the network architecture that supports these dynamics is already in place during IEDs. Furthermore, these mechanisms can be inferred through DCM and explored through simulation, combining the strength of computational modelling as in the previous study, with the constraints of empirical data recorded directly from patients.

### 4.4 Implications

Our study used TSC as an interesting test case to explore the local network constraints of interictal dynamics, because patients share an underlying aetiology, often have multiple lesions that can be recorded from during the same iEEG recording, and there are important clinical questions in relation to the involvement of perituberal cortex in ictogenesis that remain to be addressed. However, this study was not designed to address the predictive values of our derived measures in terms of clinical outcomes, as this would need to be addressed with a larger patient cohort with heterogeneous, and long-term recorded outcomes.

It is important to note that patients with tuberous sclerosis have constant growth of new tubers and maturation (calcification) of already existing tubers (Curatolo et al., 2008; Grajkowska et al., 2010), meaning that not previously ictogenic tubers can become ictogenic over time, and that new tubers can form – leading to new epileptic foci. Thus improving our surgical approaches to limit excess morbidity is essential, as patients with TSC may undergo several epilepsy surgeries during their lifetime (Bollo et al., 2008; Fallah et al., 2015).

Although our results specifically address patients with TSC, other pharmaco-resistant epilepsies are associated with pathologies in the same molecular (mTOR) pathway (e.g. focal cortical dysplasia type 2) and share histological features, therefore our results may also be applicable to that group of patients (Meng et al., 2013; Liu et al., 2014; Marsan and Baulac, 2018).

## Acknowledgements

We are grateful to all the patients and their families, as well as to the clinical teams overlooking their care. This work is supported by the National Institute for Health Research Biomedical Research Centre at Great Ormond Street Hospital for Children NHS Foundation Trust and University College London. The project was funded by the Oakgrove Charitable Foundation (MT, RT, ST, RER), and the Wellcome Trust (RER: 209164/Z/17/Z, 106556/Z/14/Z, KJF: 088130/Z/09/Z)

